# Occupancy patterns of 208 DNA-associated proteins in a single human cell type

**DOI:** 10.1101/464800

**Authors:** E. Christopher Partridge, Surya B. Chhetri, Jeremy W. Prokop, Ryne C. Ramaker, Camden S. Jansen, Say-Tar Goh, Mark Mackiewicz, Kimberly M. Newberry, Laurel A. Brandsmeier, Sarah K. Meadows, C. Luke Messer, Andrew A. Hardigan, Emma C. Dean, Shan Jiang, Daniel Savic, Ali Mortazavi, Barbara J. Wold, Richard M. Myers, Eric M. Mendenhall

## Abstract

Genome-wide occupancy maps of transcriptional regulators are important for understanding gene regulation and its effects on diverse biological processes, but only a small fraction of the >1,600 transcription factors (TFs) encoded in the human genome has been assayed. Here we present data and analyses of ChIP-seq experiments for 208 DNA-associated proteins (DAPs) in the HepG2 hepatocellular carcinoma line, spanning nearly a quarter of its expressed TFs, transcriptional co-factors, and chromatin regulator proteins. The DAP binding profiles classify into major groups associated predominantly with promoters or enhancers, or with both. We confirm and expand the current catalog of DNA sequence motifs; 77 factors showed similar motifs to those previously described using *in vivo* and/or *in vitro* methods, and 17 yielded novel motifs. We also describe motifs corresponding to other TFs that co-enrich with the primary ChIP target. FOX family motifs are, for example, significantly enriched in ChIP-seq peaks of 37 other DAPs. We show that promoters and enhancers can be discriminated based on motif content and occupancy patterns. This large catalog reveals High Occupancy Target (HOT) regions at which many DAPs associate, although each contains motifs for only a minority of the numerous associated DAPs. These analyses provide a deeper and more complete overview of the gene regulatory networks that define this cell type.

## Introduction

Transcription factors (TFs) are DNA-binding proteins that play key roles in gene regulation [1,2]. According to the most recent census and review of putative TFs, including manual curation of DNA-binding domains in protein sequences and experimental observations of DNA binding, there are 1,639 known or likely TFs in the human genome [2]. However, other tallies [1,3], and broader definitions of proteins that associate with DNA, including transcriptional cofactors (CFs) and chromatin regulators or chromatin modifying enzymes (CRs), suggest there may be as many as 2,500 such proteins encoded in the human reference assembly; we refer to these collectively as DNA-associated proteins (DAPs), in order to distinguish this broad group of proteins from the stricter definition of direct DNA-binding TFs. A typical TF binds preferentially to a short DNA sequence motif, and, *in vivo*, some TFs also exhibit additional chromosomal occupancy mediated by their interactions with other DAPs [4-6], although the extent and biological significance of most secondary associations are not well understood [7]. TFs, CFs, and CRs all play vital roles in orchestrating cell type- and cell state-specific gene regulation, including the temporal coordination of gene expression in developmental processes, environmental responses, and disease states [8-14].

Identifying genomic regions with which a TF is physically associated, commonly referred to as TF binding sites (TFBSs), is an important step toward understanding its biological roles. The most common genome-wide assay for identifying TFBSs is chromatin immunoprecipitation followed by high-throughput sequencing (ChIP-seq) [15-17]. In addition to highlighting potentially active regulatory DNA elements by direct measurement, ChIP-seq data can define specific DNA sequence motifs that can be used, often in conjunction with expression data and chromatin accessibility maps, to infer likely binding events in other cellular contexts without direct assays. Elegant methods have been developed for identifying motifs [18-21], including ones that consider the plasticity of individual bases within and adjacent to a motif [22-25], account for structural details in relation to TF co-occurrence [26-28], or incorporate directly measured and inferred motifs [4]. Subsets of motifs can be specific to different cell types or environmental contexts, and can depend on chromatin status and presence of cofactors for accessibility [29,30], and motif sequence alone is not always predictive of binding events [31-33]. While motifs identified by enrichment in ChIP-seq are often representative of direct binding, this is not always the case, as co-occurrence of other DAPs could lead to the enrichment of their motifs. Further, the ChIP-seq method identifies both protein:DNA and, indirectly, protein:protein interactions, such that indirect and even long-distance interactions (e.g. looping of distal elements) are captured as ChIP-seq enrichments.

A long-term goal for the field is comprehensive mapping of all DAPs in all cell types, but a compelling and more immediate aspiration is to create a deep map of all DAPs expressed in a single cell type. The resulting consolidation of hundreds of genome-wide maps for a single cellular context promises insights into TF/CF/CR networks that are presently not possible. It will also provide the necessary backdrop for understanding large-scale functional element assays, and should improve the ability to infer TFBSs in other cell types that are less amenable to direct measurements.

Previous analyses of sets of numerous DAPs have been performed [34-38]. However, the larger studies to date have assayed occupancy by transfected DAPs, often expressed ectopically and at non-physiological levels, in contrast to this study, in which we performed assays on endogenous proteins expressed at physiological levels. This work in the HepG2 hepatocellular carcinoma cell line is part of the Encyclopedia of DNA Elements (ENCODE) Consortium effort toward achieving “factor completeness” (e.g., the mapping of all expressed DAPs’ binding locations) in a subset of commonly used human cell lines. We present here an analysis of 208 DAP occupancy maps in HepG2, composed of 92 traditional ChIP-seq experiments with factor-specific antibodies and 116 CETCh-seq (CRISPR epitope tagging ChIP-seq) experiments. The CETCh-seq method was developed to address the dearth of ChIP-competent antibodies for many factors, and has been shown to be a robust, powerful assay [39,40]. Its strength is that the endogenous DAPs are tagged with a universal epitope that is recognized by a single well-characterized ChIP antibody, and that the tagged factors are expressed at physiological levels to avoid ectopic ChIP peaks that can be caused by conventional transgene overexpression [41,42]. As more CETCh-seq experiments are performed, the growing database is used to identify any antibody-specific artifacts attributable to cross-reactivity. This is part of the ENCODE Consortium quality control process for ChIP-seq, CETCh-seq, and related assays [43], which includes immune reagent validation and characterization by assays such as western blots, and validation of tagged cell lines by confirmation of genomic DNA sequence. Additionally, the hundreds of ChIP experiments performed have led to tuning and optimization of protocols in efforts to alleviate technical biases [44,45]. Results of validation experiments for all DAPs assayed here are available on the ENCODE web portal, at www.encodeproject.org.

Of the >1,600 total human DAPs, approximately 960 are expressed in HepG2 cells above a threshold RNA value of 1 FPKM (Fragments Per Kilobase of transcript per Million mapped reads), the minimum level at which we have obtained successful ChIP-seq and CETCh-seq results. The resource we present here contains ChIP-seq and CETCh-seq maps for ~22% of these 960 factors, of which 171 are sequence-specific TFs and 37 are chromatin regulators and transcription cofactors (Figure 1A and Supplementary Table 1). This large and unbiased sampling in one cell type allowed us to approach analysis from complementary directions, beginning with patterns of DAP occupancy and co-occupancy to find preferential associations with each other and with promoters, enhancers, or insulator functions, and in the other direction, working from genomic loci, sequence motifs, and epigenomic state to explain occupancy.

**Figure 1.**
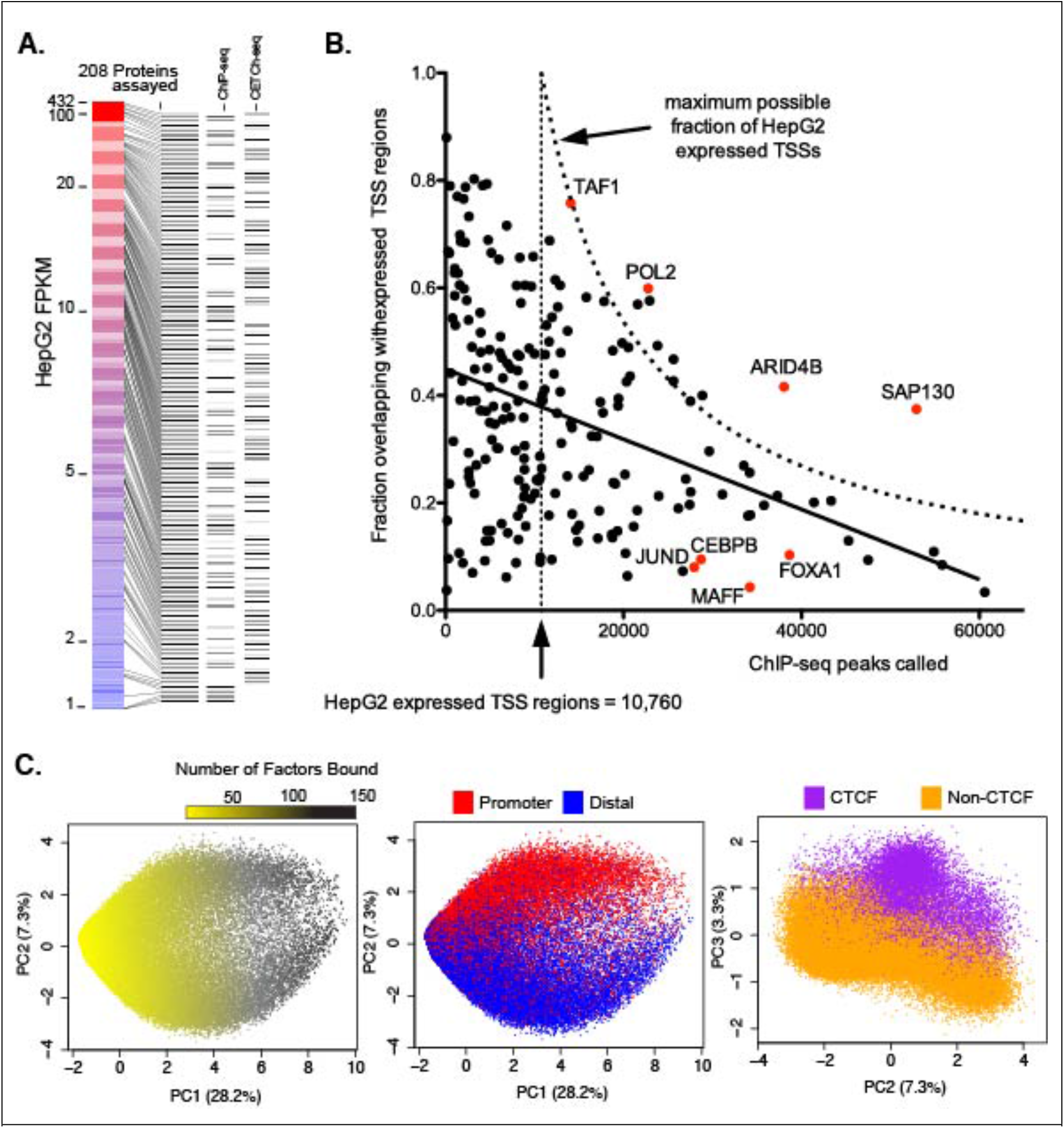
Overview and analysis of HepG2 datasets

**A.** The 208 DNA-associated factors assayed in HepG2, organized by expression (FPKM), and denoting whether the factors were assayed by ChIP-seq and/or CETCh-seq. **B.** Scatter plot of all 208 factors showing broad distribution of fraction of called peaks at expressed TSSs (+/- 3 kb of TSS) vs. total peak number; points beyond maximum possible fraction represent multiple peaks at single TSS regions. **C.** Plots showing PCA of genomic segments with more than two factors bound, highlighting the separation based on number of factors bound, promoter vs. distal, or the presence of CTCF.

All ChIP-seq/CETCh-seq data are available through the ENCODE web portal (www.encodeproject.org), as well as at Gene Expression Omnibus. Each DAP’s genome-wide binding sites were identified using the SPP algorithm [46], with replicate consistency and peak ranking determined by Irreproducible Discovery Rate (IDR) [47]. This publicly available ENCODE occupancy data, attaining the greatest factor depths at physiologically-relevant expression levels to date, together with analyses and insights presented here, comprise a key resource for the scientific community.

## Results

### DNA-associated proteins segregate underlying element types and states

As an initial analysis, we asked how the binding of each of the 208 DAPs is distributed in the genome relative to known transcriptional promoters. Specifically, we calculated the fraction of called peaks within 3 kilobases (+/- 3 kb) of transcription start sites (TSSs) for each factor, analyzing only TSSs of genes expressed (>=1 TPM, or Transcripts Per Kilobase Million) in HepG2 (Figure 1B) and, separately, all annotated TSSs regardless of expression (Supplementary Figure 1).

To further summarize the occupancy landscape, we merged all the called peaks from every experiment into non-overlapping 2 kb windows, limited to those windows in which two or more DAPs had a called peak, and performed a Principal Component Analysis (PCA) on these DNA segments, using presence/absence of each DAP at each segment. This analysis captured global patterns of ChIP-seq peaks, with Principal Component 1 (PC1) explaining ~28% of the variance and correlating strongly with the number of unique DAPs associated with a given genomic region (Figure 1C). PC2 separates promoter-proximal from promoter-distal peaks, underscoring the relevance of promoters as a major predictor of genomic state and DAP occupancy. Interestingly, the shape of this plot suggests that as the number of DAPs associated at a locus increases, the promoter-proximal and promoter-distal regions lose separation along PC2. Additionally, PC2 plotted against PC3 shows strong segregation based on occupancy of the factor CTCF (Figure 1C), suggesting discrete genomic demarcations attributable to this important factor, as expected for its insulator/loop anchoring functions.

To assess the epigenomic context of each binding site, we used IDEAS (an Integrative and Discriminative Epigenome Annotation System), a machine learning method for biochemical mark-based genomic segmentation [48]. This IDEAS HepG2 epigenomic segmentation inferred 36 genomic states based on eight histone modifications, RNA polymerase ChIP-seq, CTCF ChIP-seq, and DNA accessibility datasets (DNase and FAIRE). Importantly, IDEAS states for HepG2 were classified using mainly histone marks, augmented by only two DNA-associated ChIP-seq maps included in our dataset (CTCF and RNA polymerase). Thus, our analyses using IDEAS segmentation are not circular, as they would be if the segmentation had used all or mostly TF binding data as input. These segregate the anticipated major classes of correlations between epigenomic states in the IDEAS segmentation and DAP associations, such as enrichment of H3K4me3 at annotated promoters and H3K27ac at candidate active enhancers, as well as open chromatin status as assayed by DNA accessibility experiments, typical of TF-bound DNA. As expected, the resulting IDEAS states classified only a minority of the HepG2 genome as potential cis-regulatory elements (Supplementary Figure 2).

Clustering of DAP peak calls by the IDEAS segments of these genomic loci delineated several clear bins of genomic state associations. Specifically, we found a subset of DAPs that are preferentially associated with promoters, another subset associated with candidate active enhancers, and a third group distributed across both proximal promoter regions and likely enhancers (Figure 2A). We also found two smaller DAP-associated clusters: one associated with heterochromatin/repressed marks (including BMI1 and EZH2, both part of the polycomb repressor complex), and one with CTCF regions (including CTCF and known cohesin complex proteins RAD21 and SMC3) (Figure 2A, Supplementary Table 2). These distinct categories contain members of different classes of DAPs, and point to distinct gene regulatory pathways. Additionally, a PCA based on these IDEAS states clearly segregated the DAPs into bins that recapitulate these clusters (Supplementary Figure 3).

**Figure 2.**
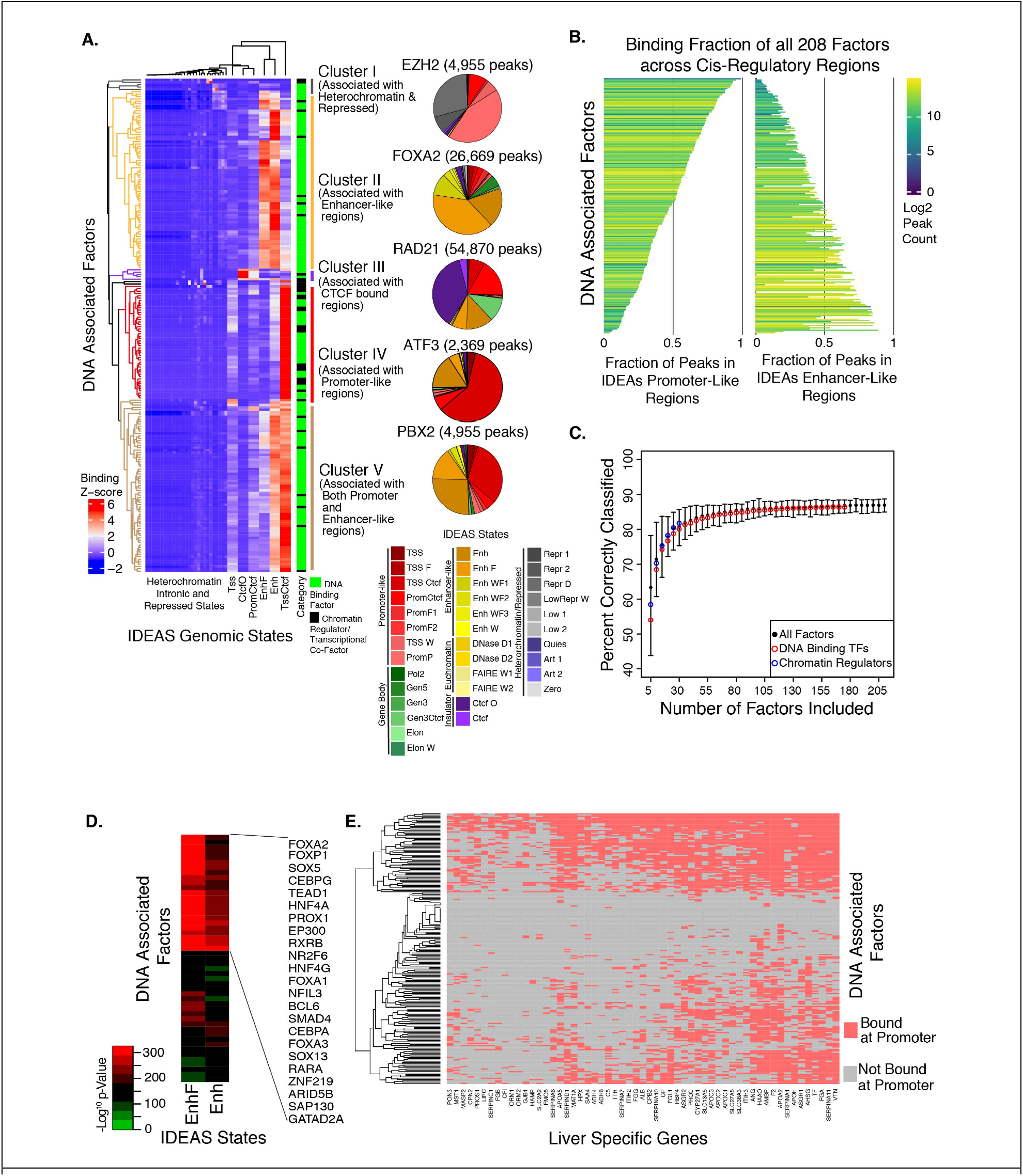
Landscape of factor binding to regulatory states. **A.** Unsupervised clustering of the 208 factors based on the binding enrichment at 36 IDEAS genome states and the 5 main clusters of factors, along with pie charts showing absolute binding fractions of an example of a factor from each cluster. **B.** Plot showing the fraction of promoter or enhancer binding for all 208 factors, with bars colored based on peak counts for each factor. **C.** Predictive ability of random forest classification of genomic regions as either enhancer or promoters based on number of factors used to train the algorithm. **D.** Enrichment of TFs at regions of the genome we classified as putative HepG2-specific cis-regulatory elements. **E.** Binding of TFs to liver specific gene promoters.

For roughly 40% of the DAPs assayed, most called peaks were in IDEAS promoter-like regions, while ~30% of DAPs were predominantly associated with IDEAS enhancer-like regions (Figure 2B). There was no significant correlation between experimental peak counts and the distribution of peaks across promoters and enhancers. While these preferences are part of a continuous distribution, the unsupervised clustering using all IDEAS genomic states suggests strong localization preferences among subsets of DAPs. The three largest subsets reveal that many DAPs are strongly enriched for promoters, while others are strongly associated with candidate enhancers, implying separable functions for the two classes of most differentiable factors. The third group in the continuum shows little or no bias, associating more equally with both promoters and enhancers. Previous publications have noted the similarities between promoters and enhancers, ascribing enhancer activity to promoters, and it is established that transcription occurs directly at enhancers in the form of enhancer-RNA (eRNA) and even as alternative promoters [49,50] (and reviewed in [51]). The subset of DAPs identified as associating with both promoters and enhancers may point to specific genomic loci or gene regulatory networks where the lines between promoters and enhancers are most blurred. It is also possible that the factors in this group are most associated with looping between promoters and distal enhancer elements. Because DAPs localize to specific genomic states, we were able to reproducibly train random forest models capable of predicting the IDEAS state of a genomic region using binding information of only a small number of DAPs (Figure 2C). The prediction method was successful when using the combination of TFs/CFs/CRs, and also when trained only on direct DNA-binding proteins or only on CFs/CRs, requiring a subset of any of ~30 DAPs to achieve ~80% accuracy.

### Liver-specific TFs and genes reveal the cis- and trans-networks of HepG2

Identifying transcription networks is important for understanding how genes specify a cell type and execute its activities. Our current understanding is that TFs, including key cell-type specifying factors, interact with other factors via combinatorial cross-regulation to drive gene expression in a cell-specific manner. To identify HepG2-specific cis-regulatory elements, we used IDEAS segmentation to identify all promoter-like and enhancer-like regions in at least one of five other cell lines (GM12878, H1hESC, HUVEC, HeLa-S3, and K562), and filtered these regions from the HepG2 segmentation. In the resulting set of 59,115 putative HepG2-specific cis-regulatory regions, we found significant enrichment (Fisher’s exact test, adjusted p-value <0.001, BH FDR corrected) of distinctive DAPs at HepG2-specific enhancer loci, including known liver-specific TFs such as HNF4A, HNF4G, CEBPA, and FOXA1, along with additional DAPs not previously associated with liver cell identity such as TEAD1, RXRB, and NFIL3 (Figure 2D).

Because HepG2 is a cancer cell line derived from liver tissue, we focused next on liver-specific genes, filtering for genes that are highly and specifically expressed in liver and also expressed in HepG2 at levels of at least 10 TPM. This identified a total of 57 key liver/HepG2 specific genes. We then examined the peak calls of all 208 DAPs close to promoter regions of the 57 liver specific genes (+/- 2 kb from TSSs), finding between 13 and 148 proteins associated with promoters of these genes. Pioneer TFs (capable of binding closed chromatin and usually involved in recruiting other factors [52,53]) such as FOXA1, FOXA2, and CEBPA, as well as key chromatin regulators such as EP300, associate with most of the 57 liver-specific genes (Figure 2E). Of note, the promoters of the very highly expressed liver genes *ALB*, *APOA2*, *AHSG*, *FGA*, and *F2* (also known as thrombin) have very high apparent factor occupancy/association: 65, 148, 124, 114, and 130 DAPs, respectively (Figure 2E, Supplementary Figure 4). We examined DAP occupancy at the promoters of all genes as well as of those genes expressed at 10 TPM or higher in HepG2, and compared these to DAP occupancy at the 57 liver-specific genes (Supplementary Figure 5, Supplementary Table 3). In each analysis, increasing factor number correlates positively with increasing RNA level. We note that some prior studies have suggested that high TF occupancy at highly expressed loci is a technical artifact of ChIP-seq [54], but, as described below in the section on HOT sites, several lines of evidence argue that these signals represent true biology. The 57 liver-specific genes have significantly higher expression (rank percentile t-test; p-value < 0.0001) when compared to other genes matched by number of DAPs, indicating a trend toward higher expression associated not only with a higher number of associated DAPs but with specific factor identities. We expanded our analysis to all genes that have higher expression than expected based on the number of DAPs associated at their promoters, identifying the particular factors enriched near these genes. For each of these DAPs, we then filtered all genes with ChIP-seq peaks called for the particular factor, ranking the expression of those genes against that of other genes with near-equal number of associated factors (within 5% of the number of associated factors). We identified DAPs that are associated with higher than expected expression, including unsurprising factors such as PAF1 and RNA polymerase II subunit A (Ser2 phosphorylated), marks of active transcription, as well as ATF4 and HSF1 (Supplementary Figure 5). However, we note that there are still many DAPs that have not yet been assayed by ChIP-seq, and this could explain some of the deviation from expected expression.

### Distribution of DNA-associated proteins in putative cis-regulatory elements

Though the 208 factors do not represent a complete catalog of all expressed factors in HepG2, we asked how much of the regulation in this cell line is captured by this partial compendium. We used IDEAS to define a set of 370,570 putative HepG2 cis-regulatory elements classified as promoters, “strong” enhancers, or “weak” enhancers (according to standard segmentation terminology). Discrete regions were specified by the IDEAS genomic segmentation, and were cataloged independent of their individual sizes, with merging of similar features within 100 base pairs (bp). This resulted in a broad size distribution, ranging from 200 bp to 12-16 kb; the larger segments usually represented locus control regions, divergent promoters (large, bidirectional promoters), or other similar significantly large genomic features (Supplementary Figure 6). We then calculated how many DAPs were associated in each of the 370,570 regions (Supplementary Figure 6). In terms of the general distribution of DAPs across all putative regulatory regions with called peaks, there are on average seven DAPs associated at any region, while 18% of the regions have only 1-5 called DAPs. Approximately 67% of the chromatin regions do not contain any called peaks; however, the vast majority of these (~85.5%) are classified as “weak” or “poised” enhancers by the IDEAS segmentation, and this class of elements is most likely to have the fewest number of associated factors and would therefore be more sensitive to completeness of assayed factors. It is also possible that these elements have undetectable levels of DAP occupancy or do not associate with any DAPs at all. Conversely, elements classified as promoters and “strong” enhancers by IDEAS are enriched for occupancy by higher numbers of DAPs (Supplementary Figure 6). Of the IDEAS-determined active promoter-like regions in the HepG2 genome, 61% contain a called peak for at least one DAP in this dataset, and of the “strong” enhancer-like regions, 75% contain at least one called peak. This analysis shows that the majority of promoters and “strong” IDEAS-modeled enhancers have one or more DAPs associated, and that these occupied elements display an unexpectedly high average of 15 and 18 called per region, respectively.

Thus, these data capture a substantial overview of the TF/CF/CR regulatory network in HepG2 cells.

### Motif analysis reveals direct binding targets and factor associations

We assessed motif enrichment in peaks, and found many previously derived motifs for both direct and potentially indirect associations, as well as a small number of potentially novel motifs. To derive and map motifs for each factor, we used the MEME software suite, TOMTOM, and Centrimo [20,21,55-58] to call and assess motifs for each experiment. We focused only on motifs called from the 171 putatively direct DNA-binding TFs in our dataset, based on previous curation [2], filtering these motifs by significance (MEME E-value <1e-05) and enrichment (CMO E-value <1e-10) to obtain a high-confidence set of 293 motifs called from 160 TFs. We compared these motifs to the JASPAR databases [59,60] and to the CIS-BP database [4] to determine whether our *de novo* derived motifs matched previous findings from various *in vivo* and/or *in vitro* assays [61]. Overall, >80% of the 293 motifs had a similar motif in these databases (86% in CIS-BP build 1.02, 82% in JASPAR2018, 81% in JASPAR2016; Supplementary Figure 7). For 103 motifs derived from peaks for 77 unique TFs, the most similar motif in the database was annotated as the motif for the TF which was the target of the ChIP/CETCh-seq assay, and we term these cases “concordant” (Figure 3A, Supplementary table 4). There were 163 motifs derived from peak data for 103 TFs that were more similar to the database motif of a different TF, and we denote these as “discordant”. We also observed 27 motifs derived from peaks of 17 TFs that were highly dissimilar to any motifs in the databases and may be novel motifs; most of these were from Zinc-Finger TFs, a large class of factors that is virtually unassayed by endogenous ChIP-seq.

**Figure 3.**
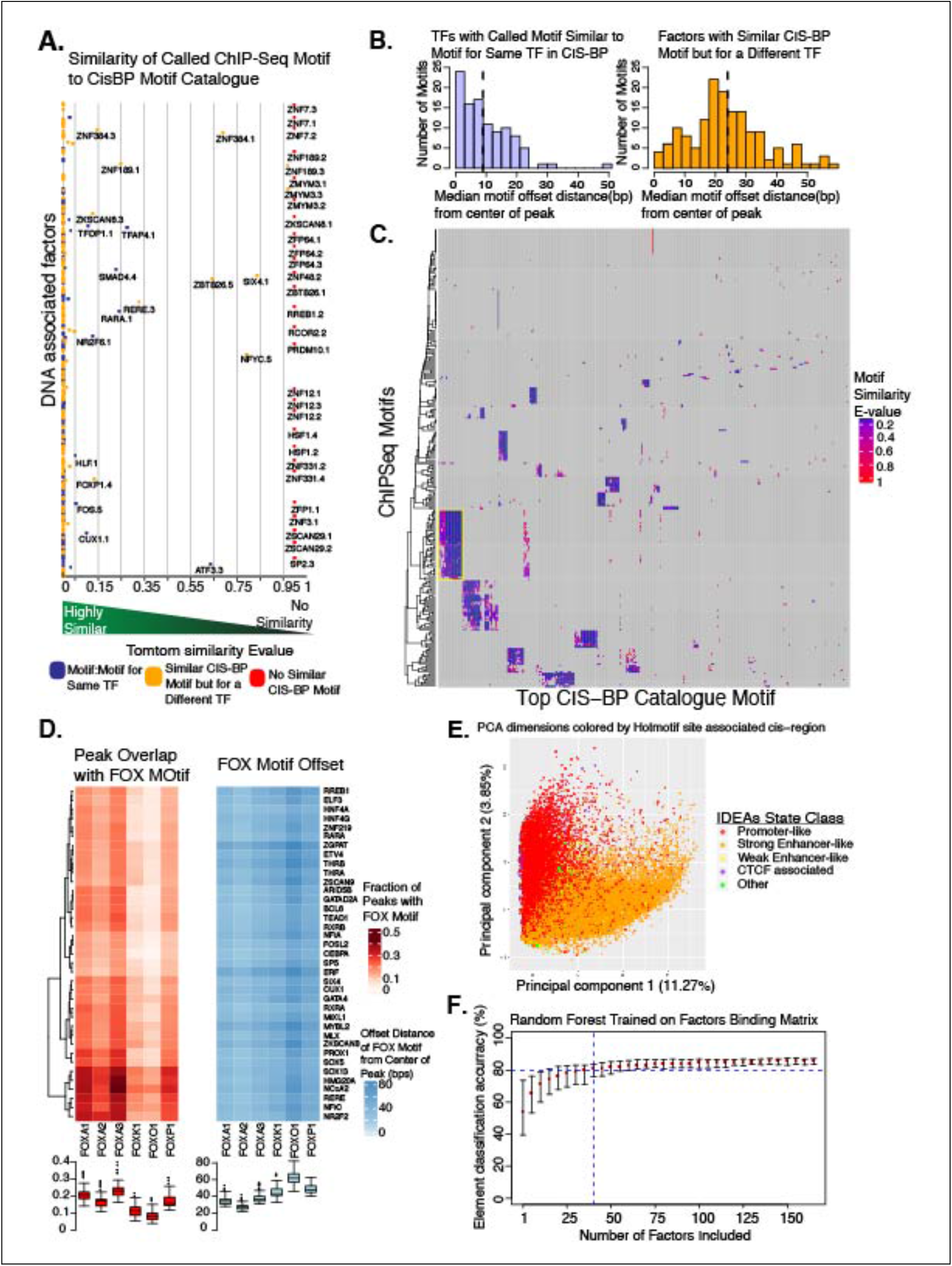
Motif Identification and Analysis. **A.** The 293 high-confidence motifs derived from analysis of the ChIP-seq data were quantitatively compared to all (human) motifs in the CIS-BP database and plotted based on similarity scores. Blue points represent motifs that matched the assayed factor, yellow points represent motifs that match a factor other than the one assayed, and red points represent motifs not similar to any in CIS-BP. **B.** Histograms showing the distance from the center of the ChIP-seq peak for motifs that match the TF, and for motifs that do not match the TF. **C.** Clustered heat map showing the similarity of all 293 significant motifs to 733 motifs from CIS-BP for the assayed factors. **D.** Further analysis of the cluster containing 37 factors that had FOX family motifs, showing the overlap of FOX TF binding in these peaks, as well as the median offset of the FOX motif from center of the ChIP-seq peaks. **E.** PCA showing separation of motifs that fall in promoters vs. those that fall in enhancers. **F.** Prediction accuracy for calling whether an element is a promoter or enhancer based on motifs present.

Examining the 163 discordant motifs, we observed an enrichment of motifs representing pioneer TFs such as FOXA1, and we hypothesize that these motifs were called due to their significant co-occurrence with the assayed TFs. Previous studies have noted the enrichment in ChIP-seq data of sequences that do not appear to be binding motifs for assayed TFs, but rather are more similar to other TF motifs [62]. There are multiple potential explanations for why the ChIP-seq derived motif would most closely match a motif previously annotated for another factor. Related TFs often recognize very similar sequence motifs; for example, the motif we derived for TEAD4 was very similar to the motif previously found for TEAD1 [63]. There are also instances where a factor lacks a strong and specific DNA binding domain and no motif would be expected unless the motif represents a frequent co-binding partner, a scenario we explore below with GATAD2A, and also seen with HMG factors. A similar explanation involves a particular TF acting as an “anchor” at a locus, and through either direct protein:protein interactions, or by inducing an open chromatin environment, behaves as the mechanism for localization of other proteins to that region of DNA. A well-studied example of this highlighted in our data was the enrichment of the CTCF motif in RAD21 ChIP-seq, as RAD21 lacks a DNA-binding domain but is known to interact with CTCF. It is difficult to confidently determine whether a discordant motif represents a key co-factor interaction or a commonly co-localized protein. We note that when we called multiple, distinct, high-confidence motifs in a single ChIP-seq experiment, with one motif annotated in databases as the direct target of the assayed TF and another motif representing a different TF that we also assayed separately, we were able to observe from the secondary factor’s ChIP-seq experiment that both TFs are likely associated at these loci, since both experiments yielded called peaks at these loci.

Supporting our hypothesis that the secondary factor’s motif was not a site of direct binding for the primary factor, an examination of the precise location of the motifs within peaks showed a significant difference (K-S test p-value < 2.2e-16) where the direct matching motifs of the assayed factors are closer to the center of called peaks, and the discordant motifs for other TFs are more offset, providing evidence for co-occurrence at these locations (Figure 3B). Direct interaction and co-recruitment between these pairs of TFs could explain these observations, and numerous examples of such combinatory and cooperative activities between TF pairs have been reported (reviewed in [64]). We also found no significant trend for secondary TF motifs in any factor clusters we identified by IDEAS state preferences or other methods, suggesting that no biases were introduced by contributions from particular genomic loci (Supplementary Figure 8). Additionally, we analyzed the peak locations of the 27 novel motifs (representing 17 factors) that were highly dissimilar to any motifs in CIS-BP, and the majority showed enrichment at the center of peaks (Supplementary Figure 9), supporting the notion that these motifs represent direct DNA binding for these factors.

To better understand discordant TF motif calls, we constructed a similarity heatmap using all 293 high-confidence motifs from our data and the motif for each assayed TF annotated in the CIS-BP database (n=733) as provided by the MEME suite software (Figure 3C). This analysis clustered TFs both by similarity of their direct binding motifs (such as all Forkhead factors) and by co-occurrence with other motifs. In this way, we were able to identify TFs that associate at genomic loci near particular motifs, such as CTCF. Most obvious was a set of 37 factors for which a Forkhead motif was called, indicating the high prevalence of this motif in HepG2 at enhancers and promoters, and the key role of factors such as FOXA1 and FOXA2 in the gene regulatory network in these cells. We examined these cases using our ChIP-seq data from six FOX TFs (FOXA1, FOXA2, FOXA3, FOXK1, FOXO1, and FOXP1), asking how often each of these FOX TFs yielded called peaks with a FOX motif that overlapped with a peak for any of these 37 other factors, and we found that most of the 37 contained a FOX peak with FOX motif in about 20% of their peaks, with FOXA1 and FOXA3 motifs being the most common (Figure 3D).

We next examined the location of the FOX motif in the overlapping peaks and found that all were offset to varying degrees, though always with median distance more than 20 bp from the center of peaks (Figure 3D). Additionally, we examined all peaks called for each of the 37 factors and identified the fraction containing a primary motif specific to the individual factor along with a FOX motif, the fraction containing only the primary motif, the fraction containing only a FOX motif, and the fraction containing neither motif (Supplementary Figure 10). For most of the 37 factors, the majority of peaks did not contain a primary motif, a result that may indicate protein:protein interactions and/or looping events in these peaks. Further, examining peak overlaps between these 37 factors and the six FOX TFs, we observed varying associations and co-occupancy partners, including factor preferences for individual FOX TFs, as well as a cluster of components of the nucleosome remodeling and histone deacetylase (NuRD) complex (Supplementary Figure 10).

We also found that motif information alone was predictive of genomic segments, clearly showing segregation between IDEAS states in a PCA (Figure 3E). A random forest algorithm trained only on motifs was able to predict IDEAS states almost as well as the method trained on ChIP-seq peaks, achieving ~80% success with any ~40 motifs (Figure 3F).

### Known and novel associations between factors

TFs and chromatin regulatory proteins can interact with and recruit other DAPs through direct and indirect physical association. While the activity of a few key TFs may be very important for cell-state expression, it is likely that combinatorial events are necessary to fine tune expression [65]. We found both known and novel associations by examining occupancy overlaps and trends in a variety of analyses.

To identify candidate co-occupancy events mediated by direct DNA binding or by indirect interactions, both of which produce peaks in ChIP-seq data, we performed several analyses. We used the PCA of the protein-bound genomic loci described above (in which genomic loci clustered according to the DAPs associated at each region; Figure 1C-E), and generated a correlation matrix based on the cumulative principal component distances (weighted by the proportion of variance explained by each component) between all DAPs. The resulting unsupervised clustering of respective pairwise distances highlighted punctate groups representing both known and potentially novel complexes, including a group containing POL2 and TSS-associated chromatin modifying enzymes, a group of cohesin complex members, a group of liver-specific factors, and a group containing the NuRD complex, among others (Figure 4A).

**Figure 4.**
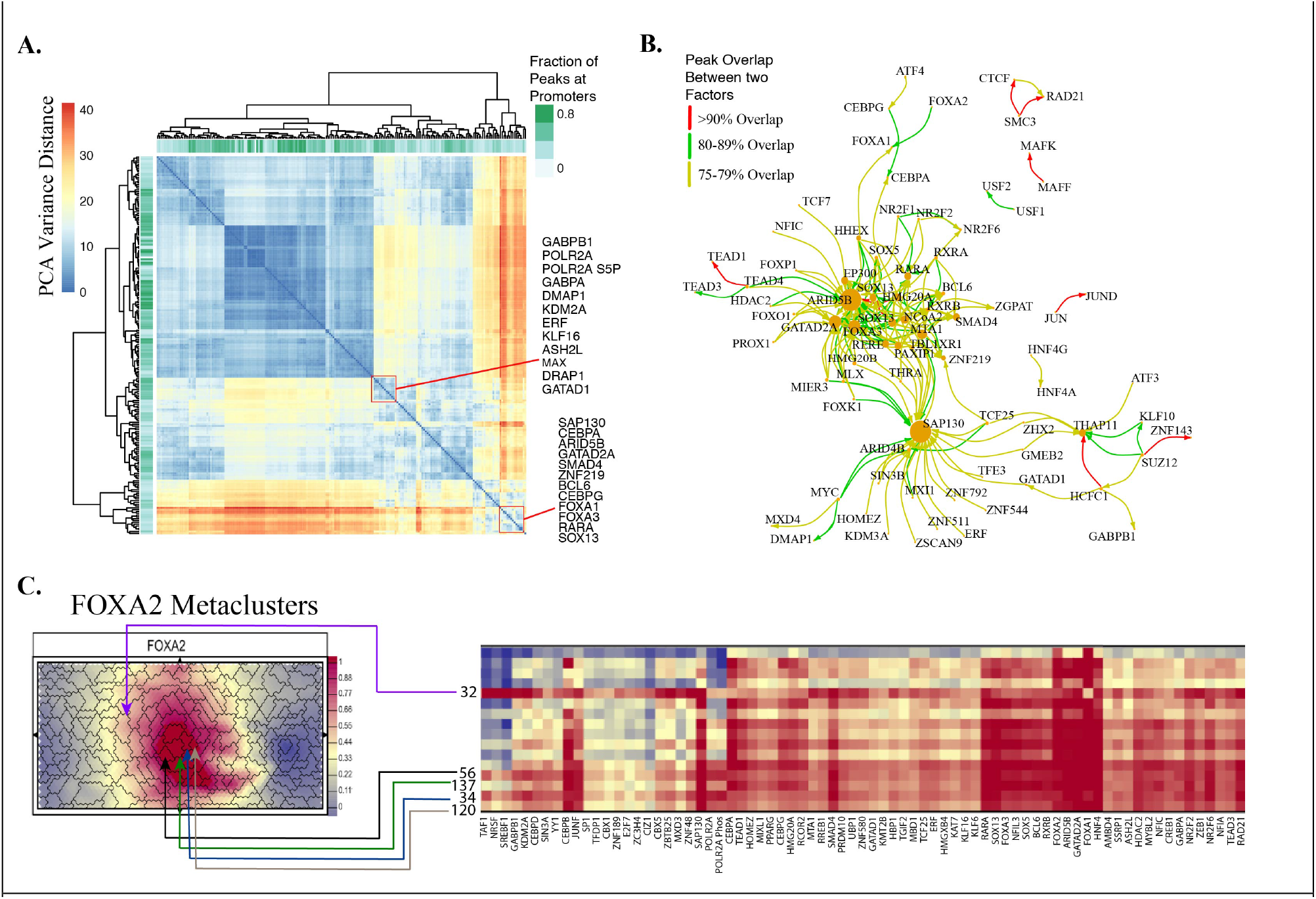
Co-localization of factors. **A.** Correlation matrix based on the cumulative principal component distances weighted by the proportion of variance explained by each component between all factors, derived from the PCA of all genomic loci with a peak containing at least two factors. **B.** Subset of network plot derived from peak overlaps between all factors showing strong associations between a subset of factors. **C.** Self-organizing map for FOXA2 in HepG2, with metaclusters showing major associations with specific factors.

We performed read count Spearman correlations between all 208 DAPs by calculating raw sequencing counts at every unique locus present in called peaks in any experiment (+/- 50 bp from peak center). The resulting correlation heatmap also showed clusters of related proteins as well as both known and potentially novel interactions (Supplementary Figure 11). Network plots based on pairwise peak overlaps highlighted a number of known interactions, including CTCF/RAD21 and CEBPA/G networks, as well as DAPs that associate with a large number of other factors, usually chromatin regulatory proteins such as SAP130, GATAD2A, and ARID5B (Figure 4B). We examined the associations at the level of called motifs by finding the peaks in each experiment where a specific called motif was present, limiting the analysis to the 293 high-confidence motifs from the 171 TFs in the data set. Upon identification of the primary motif, we looked for associations between motifs 1-40 bp away (Supplementary Figure 12). This analysis reveals the TFs (and motifs) that are more likely to associate with any other particular TF’s motif. Of note, we observed that RAD21 is highly associated with CTCF motifs, as expected, and we also found several other known complexes as well as some novel associations. We found that FOXA1 peaks with the canonical Forkhead motif are more likely to contain relatively few motifs for other factors, but that many factors, such as HNF4A, HNF4G, and RXRB, are enriched for nearby FOXA1 motifs.

For an independent assessment of co-occupancy, we trained a chromatin self-organizing map (SOM) [66] using all 208 DAPs with the SOMatic package [67]. This analysis generated 196 distinct clusters of SOM units, with each such “meta-cluster” sharing similar profiles, and corresponding decision trees that trace the supervised learning path used to determine the unique features of each metacluster profile (Figure 4C, Supplementary Figures 13, 14). Focusing on the key HepG2 transcription factors FOXA1/2 and HNF4A, we found that 18 distinct metaclusters accounted for nearly half of the peaks for these 3 TFs (43% for FOXA1, 43% for FOXA2, and 49% for HNF4A). DAPs important for liver development, nucleosome remodeling, and the cohesin complex show high co-binding signal in these key 18 metaclusters.

Looking closer at the DAPs that distinguish these 18 key clusters, we found that five of these (numbered as 32, 34, 56, 120, and 137) show strong signal from CEBPB, SAP130, and RAD21 (Figure 4C, Supplementary Figure 13). In particular, metacluster 32 had a collection of unique features related to the NuRD complex and liver processes (Supplementary Figure 13). A decision tree trained on regions in this cluster highlighted the presence of TAF1 and MTA1 (part of the NuRD complex) and the absence of a high signal of KLF16 (a known TF displacer) as sufficient to predict association with MBD1, HBP1, and HDAC2 (a sub-unit of the NuRD complex) with ~91% accuracy. GREAT (Genomic Regions Enrichment of Annotations Tool [68]) analysis of these regions revealed a related set of negative regulation and response GO terms (Supplementary Figure 13), which provides further evidence that the NuRD complex is involved in tissue specific gene regulation.

The indirect motif, co-occupancy, and SOM analyses led us to find novel factors associated with GATAD2A, a core component of the NuRD complex. GATAD2A has been recalcitrant to antibody ChIP-seq and therefore was one of the targets for our CETCh-seq protocol. The experiments revealed that 53% of the GATAD2A peaks in HepG2 are annotated as active enhancers (Figure 5A), a surprising observation given the association of the NuRD complex with transcriptional repression and enhancer decommissioning [69-71]. GATAD2A has a very degenerate DNA binding domain, and is not predicted to bind DNA independently, and indeed we found the called GATAD2A motif to match FOXA3 (Figure 5B). In our co-association analysis in HepG2, we identified 6 factors that co-occur in discrete genomic regions with GATAD2A (Figure 5C). We analyzed our GATAD2A-FLAG protein immunoprecipitation by mass spectrometry, and this revealed that multiple components of the NuRD complex also coimmunoprecipitate with GATAD2A (Supplementary Table 5). Of the GATAD2A-associated proteins, ZNF219 [72], SMAD4 [73], and RARA [74] have previously been associated with the NuRD complex (Figure 5C). We additionally identified ARID5B, FOXA3, and SOX13 as proteins associated with the known NuRD group, specifically at active enhancers with enrichment of Forkhead binding sites (Figures 5B, 5C). The classic NuRD complex has been suggested to function at enhancer regions associated with tissue-specific gene regulation [75], and our data confirms that the core NuRD component GATAD2A is recruited into these regions. Of note, NuRD binding at these open and presumably active regions is thought to function through a NuRD complex containing MBD3 and not MBD2, and our GATAD2A-FLAG IP-mass spectrometry data confirmed this, as we observed MBD3 peptides but no MBD2 peptides immunoprecipitated with GATAD2A (Supplementary Table 5) [76].

**Figure 5.**
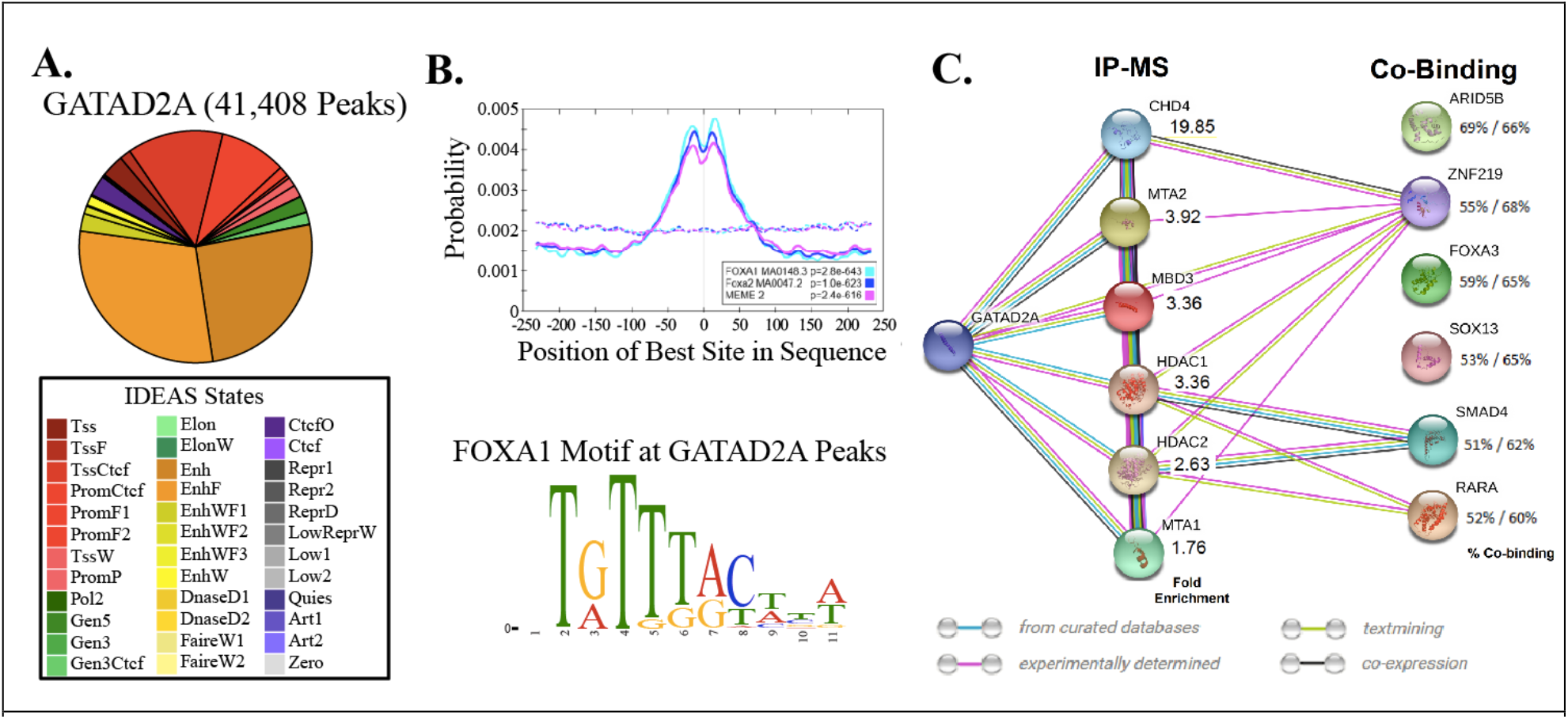
GATAD2A co-localization analysis. **A.** IDEAS state binding for GATAD2A showing enrichment at enhancers. **B.** Presence of top motifs at GATA2DA bound regions and the top motif called at these peaks. **C.** NuRD complex members and their identification through IP mass spec of GATAD2A IPs, and through co-binding at GATAD2A bound loci.

### Highly occupied regions are driven by individual TF binding

We examined how many factors were bound at each putative cis-regulatory element by merging all peaks from all 208 DAP experiments, with a maximum merged size of 2 kb. This analysis yielded a total of 282,105 genomic sites with at least one associated DAP, a mean of 7.36 associated DAPs, and maximum of 168 DAPs. We asked if certain DAPs are more likely to co-occupy at genomic loci with a high number of other DAPs. To answer this, we performed hierarchical clustering of the degree of co-association for each DAP, which results in three distinct clusters (Figure 6). The first is a cluster of 33 proteins, including previously described key pioneer factors such as FOXA1 and FOXA2 [77], which exhibit a low degree of co-occupancy with other DAPs at a relatively high proportion of their binding sites [78]. The second cluster, comprised of 32 DAPs, displays frequent association at higher co-occupancy regions and is composed of DAPs already known to be recruited by, or to interact with, a large number of other factors, such as MYC and DNMT3B [79,80]. The third cluster contains the remaining DAPs, which exhibit an intermediate degree of co-occupancy, including key HepG2 TFs such as HNF4A and FOXA3.

**Figure 6.**
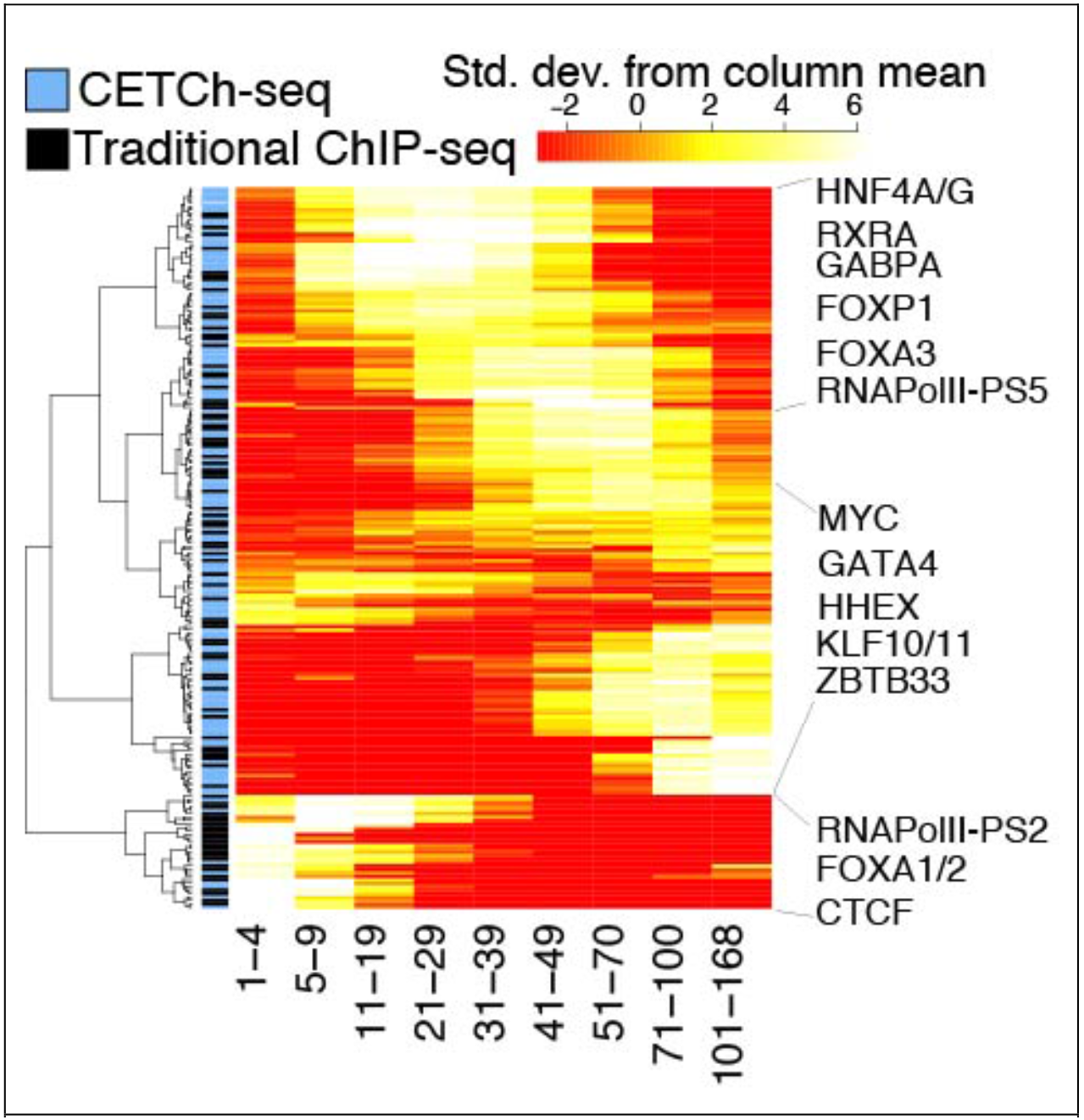
Factor enrichment at loci with increasing number of factors bound.

As previously described [81-83], there are many regions in the genome occupied by large numbers of DAPs in ChIP-seq assays (example shown in Supplementary Figure 15). There are several possibilities to explain these High Occupancy Target (HOT) regions [84]. Some researchers have filtered all or the majority of these regions from analyses under the assumption they are artifacts [54,85]. It is also possible that they are the result of stochastic shuffling of direct binding of many DAPs in a population of cells; when assayed across the millions of cells used for an individual ChIP-seq experiment, this could result in apparent co-localization of peaks for many DAPs which are not actually co-occupied at the same time in the same cell. Mechanisms underlying this might include indiscriminant recruitment driven by key factors or some unknown property of these regions of open chromatin, or by densely packed DNA sequence motifs. It is also conceivable that three-dimensional genomic interactions, including enhancer looping and/or protein complexes, lead to ChIP-seq cross-linking of DAPs in close proximity.

We define HOT regions in these data as those sites with 70 or more DAPs within a 2 kb region (n=5,676). Intersecting HOT regions with the previously described IDEAS segmentations revealed that greater than 92% of HOT regions map to candidate promoter or “strong” enhancer-like states (42.25% and 49.88% respectively). We determined using GREAT analysis that promoter-localized HOT regions are associated with housekeeping genes and that distal enhancer HOT regions are near genes associated with liver-specific pathways (Supplementary Figure 16). Additionally, we observed that higher numbers of factors in a particular locus correlates with higher expression of the nearest gene (as discussed above) and with higher sequence conservation (Supplementary Figures 17, 18). While previous researchers have noted apparent general ChIP bias favoring highly expressed genomic regions [54], we are able to perform ChIP in untagged cells with an antibody raised against the epitope tag used in CETCh-seq experiments, normalizing for this background in peak-calling, and the HOT regions continue to be strongly enriched (data not shown).

We computationally examined the general DNA motif structure of the HOT sites using PIQ (Protein Interaction Quantification) [86]. Using TF footprints identified in ENCODE HepG2 DNaseI hypersensitivity data by PIQ, we observed that at a given locus the number of TF footprints is significantly positively correlated with the number of factors that have called peaks in the locus (Supplementary Figure 19). This observation was true at multiple PIQ purity (positive predictive value) thresholds and also when using TF footprints called in the same data set from JASPAR motifs. This is consistent with HOT regions having TF motif-driven architecture as a major characteristic. To determine whether factor occupancy at highly bound regions is driven by specific DNA motifs, we trained a Support Vector Machine (SVM) on “HOT-motif” sites, a set of peaks with 50 or more co-localized motifs derived from the HOT sites (n=2,040). We tested the SVM’s predictive ability as the number of TFs increased, and observed that predictions remained constant, rather than declining, further strengthening the notion that these sites are not artifacts (Supplementary Figure 20). Precision Recall Area Under Curve (PR-AUC) scores for the SVM averaged at ~0.74 for motif-level predictions, and ~0.66 for peak-level predictions, scores substantially higher than expected, given the random sample of a positive set of 5,000 sites tested against 10X GC-matched null sequences as the negative set (Supplementary Figure 21). We also found, using the k-mers generated by the SVM, that there are 1-5 TFs at each site with very high motif affinity, and ~25-50 TFs with degenerate or weaker motifs (Supplementary Figure 22), and this observation was true when examining both HOT-motif sites and the broader HOT sites.

We asked whether this observation was unique to HOT regions (n=5,676) when compared to an equal number of enhancer regions with only 2-10 associated factors or to a null set of random enhancer elements with any number (0-208 DAPs) of associated factors (as defined by IDEAS segmentation). We observed that the sites with 2-10 factors had significantly fewer numbers of both high-affinity and low-affinity TF motifs, and that the random enhancers were essentially devoid of strong motifs (Supplementary Figures 22, 23). Indeed, the distribution of SVM scores in HOT sites was significantly higher than that of the SVM scores of sites with 2-10 associated factors, and both were significantly higher than that of the null set of random enhancer elements, indicating that the information imparted by the DNA sequence of HOT sites exceeds that of other cis-regulatory elements (Supplementary Figure 24). Moreover, in HOT sites, the strongest affinity TF at any individual peak varied across sites, indicating regulatory roles attributable to many different factors. The analysis identified important liver factors, such as FOXA3, HNF1A, and CEBPA exhibiting the strongest putative motif affinity at many of these sites (Supplementary Figure 25). This supports the notion that HOT sites are driven by a few strong and specific TF-DNA interactions and non-specific recruitment of other factors, likely through both protein complexes and binding to degenerate motifs, and possibly linking together multiple distal genomic regions through DAP interactions. This further justifies the importance of generating complete DAP maps to determine the full complement of DAPs associated at each locus, an outcome that would not occur by analysis of functional motifs only.

## Discussion

This study introduces a community data resource of occupancy maps for human transcription factors, transcriptional co-factors, and chromatin regulators that illustrates the strengths of building toward a complete catalog of DAP interactions in an individual cell type. At this intermediate stage of factor-completeness (~22% of all expressed DAPs in HepG2) the aggregated data enabled us to identify multiple known complexes and associations through various analyses, and to identify putative novel associations for future research. We also gained new insights into gene regulatory principles, clearly showing the segregation of categories of factors associated with varying localization at particular genomic states.

We approached our analysis from complementary directions, analyzing occupancy from the perspective of factor occupancy patterns and from the perspective of genomic loci and the factors that associate at those sites. Multiple analyses showed that some DAPs, including TFs, associate preferentially at promoters, while others, including different TFs, prefer enhancers. They are parts of a continuous distribution, and many factors are associated with both proximal and distal elements in varying degrees. This broad gradient of function among DAPs now poses questions about the underlying mechanisms.

The large number of factors assayed provided the capacity to identify and study regions of the genome associated with very high numbers of DAPs, compared with expectations from detailed work on specific enhancer complexes like the interferon enhanceosome [87]. Multiple lines of evidence argue that, as a group, the regions with high numbers of factors detected are neither biological noise associated with general open chromatin nor ChIP-seq/CETCh-seq technical artifacts. HOT regions have been previously described as being depleted of TF motifs, but we now suggest that this was likely due to the fact that earlier analyses lacked a large enough sampling of key TFs with strong “anchoring” motifs. Our current analyses were informed by a much larger sampling of TFs and other DAPs, and they lead us to propose a model in which HOT regions are nucleated by anchoring DNA motifs and their cognate TFs. They would form a core, with which many other DAPs can and do associate by presumed protein:protein interactions, protein:RNA interactions, and relatively weak DNA interactions at poorer sequence-motif matches.

Extensive apparent co-occupancy at domains possessing few or zero anchor motifs can potentially be explained when the ChIP assay captures, through presumed protein:protein fixation, non-adjacent DNA regions that associate with each other by looping interactions.

It is important to appreciate that the standard ChIP assay is performed on large cell populations. This means that patterns of computational co-occupancy, which we report on here, cannot discriminate between the simultaneous association of many factors in a single large molecular complex versus diversified smaller complexes that are distributed at any given time across the cell population, with each containing a smaller number of secondary associations, that sum to give massive computational co-occupancy. We can, however, state that at individual known transcriptional enhancers with >70 factors, the ChIP signal for identified anchor factors was significantly higher in magnitude.

The results thus far argue that a fully comprehensive catalog of all DAPs will help us to parse among these possibilities, which are not mutually exclusive. Completeness should also contribute to identification of additional novel motifs, and, in the cases of indirect motifs found for factors with known direct motifs, allow for more accurate motif-calling. Additionally, a complete catalog of factors in a single cell type will support imputation of critical contacts in DAP networks for three dimensional assembly of genomic enhancer-promoter organization not possible from a few individual DAP binding maps, as demonstrated by our findings regarding the NuRD complex.

We anticipate the continued addition of data from more DAPs, and aim to achieve factor completeness in at least one cell line, and hopefully more. We are very interested in learning which of the patterns we observe are specific to HepG2, and which will be recapitulated in other cell lines and, importantly, in primary cells or tissues. The ENCODE Project also continues to expand cellular contexts for these assays. We anticipate more large-scale analyses such as this, and hope that the perspectives gained from these inform more targeted research endeavors and generate meaningful hypotheses.

## Methods

### Data access

Data sets generated from this study are available at the ENCODE portal and at Gene Expression Omnibus (https://www.ncbi.nlm.nih.gov/geo/) under accession number GSE104247

### ChIP-seq/CETCh-seq

All protocols for ChIP-seq and CETCh-seq are previously published and available at the ENCODE web portal (www.encodeproject.org/documents) [17,52]. Briefly, pools of cells were grown separately to represent replicate experiments. Crosslinking of cells was performed with 1% formaldehyde for 10 minutes at room temperature and the chromatin was sheared using a Bioruptor^®^ Twin instrument (Diagenode). Antibody Characterization Standards are published on the ENCODE web portal and consist of a primary validation (western blot or IP-western blot) and a secondary validation (IP followed by mass spectrometry) for traditional antibody ChIP-seq. With CETCh-seq experiments, a molecular validation (PCR or Sanger sequencing confirmation of edited genes) in addition to one of the immunological validations (western blot, IP-western blot, or IP-mass spectrometry) is required for release. Raw fastq data were downloaded from the publicly available ENCODE Data Coordination Center, and aligned to human reference genome (hg19) using BWA-0.7.12 (Burrows Wheeler Aligner) alignment algorithm [88]. Post alignment filtering steps were carried out by samtools-1.3 [89] with MAPQ threshold of 30, and duplicate removal was performed using picard-tools-1.88 [http://picard.sourceforge.net]. Followed by filtering, each TF’s genome-wide binding sites (peak enrichment) were computed using phantompeakqualtools, implementing SPP algorithm [43,46], with replicate consistency and peak ranking determined by Irreproducible Discovery Rate (IDR) using the IDR-2.0.2 tool [56] to generate narrowpeaks passing IDR cutoff 0.02 (soft-idr-threshold). ENCODE blacklisted regions (wgEncodeDacMapabilityConsensusExcludable.bed.gz, downloadable from UCSC genome browser https://genome.ucsc.edu/) were filtered out. Additionally, we note that plasmids used to generate edited cells with epitope-tagged TFs are deposited to

Addgene, the non-profit plasmid repository, and are available for researchers to tag these factors in other cell lines of interest. We also note that GC content of DNA has been reported as a source of bias in ChIP-seq data, leading to over-representation of TFBSs and false positive peak calls, which could confound subsequent analyses [90,91]. To address this concern, we have performed ChIP-seq experiments in unedited cell lines using the FLAG antibody (Sigma F1804) utilized in CETCh-seq, and used these libraries as background for peak-calling. In these experiments, the only variable is the edited cell line used as foreground, and most biases should be accounted for.

### De novo sequence motif analysis

To identify enriched sequence motifs in the binding sites of sequence-specific factors, de novo sequence motif and motif enrichment analysis was performed using MEME-ChIP [56] suite and pipeline was built as previously described [57], on 500 bp regions centered on peak summits based on hg19 reference genome fasta. Top 5 motifs per dataset were reported from top 500 peaks based on signal value, using 2X random/null sequence with matched size, GC content and repeat fraction as a background. Central motif enrichment analysis was performed using Centrimo [21], to infer most centrally enriched motifs with de novo motifs generated from the pipeline against the 2X null sequence background.

### Comparative motif analysis

De novo motifs generated from DNA binding factors were filtered for high confidence motifs, including only highly significant and strongly enriched in binding sites, based on MEME E-value < 1e-05, Centrimo E-value < 1e-10 and Centrimo binwidth < 150. High confidence motifs were then compared, and quantified for similarity against the previously derived or known motifs available in the CIS-BP build 1.02 and JASPAR 2016/2018 databases [4,59,60] using TOMTOM quantification tool [58]. TOMTOM E-values < 0.05 represent highly similar motifs, and > 0.05 represent the motifs with increasing magnitude of dissimilarity, or more distantly related motifs.

### Gene expression

RNA-Seq quantification data (TPM, transcripts per million) for 56 cell lines and 37 tissues were retrieved from Human Protein Atlas (version 17, downloadable from https://www.proteinatlas.org/) [92], and used to identify 57 genes highly and specifically expressed in liver as compared to all other cell and tissue types, and also found in HepG2 with at least 10 TPM. On average, these 57 liver specific genes were 151.21 times expressed than any other cell types.

### IDEAS segmentation

IDEAS segmentation for six cell-types – HepG2, GM12878, H1hESC, HUVEC, HeLaS3, and K562 – were collected from the Penn State Genome Browser (http://main.genome-browser.bx.psu.edu/). All promoter-like and enhancer-like regions identified in at least one of five other cell lines, were merged using pybedtools [93,94] and these regions were filtered from the HepG2 segmentation. Significant enrichment of TF’s in the cis-regulatory regions was evaluated using Fisher’s exact test (pval adjusted<0.001, BH FDR corrected) against random or null sequence with matched length, GC content and repeat fraction using null sequence python script from Kmer-SVM [95]. Heatmaps were generated using heatmap.2 function from R gplots package [https://cran.r-project.org/web/packages/gplots/].

### GREAT analysis

Cis-regulatory associated highly TF bound sites were binned into promoter-associated and enhancer-associated sites using IDEAS segmentation. To assess the biological function and relevance of these highly TF occupied sites, GREAT (Genomic Regions Enrichment of Annotations Tool) [68] analysis was performed to predict the function of TF bound cis-regulatory regions (http://bejerano.stanford.edu/great/public/html/) associating the genomic regions to genes from various ontologies such as GO molecular function, MSigDB and BioCyc pathway. The parameters used for GREAT analysis were Basal+extension (constitutive 5.0 kb upstream and 1.0 kb downstream, up to 50.0 kb max extension) for all enhancer-associated sites, and Basal+extension (constitutive 5.0 kb upstream and 1.0 kb downstream, up to 5.0 kb max extension) for all promoter-associated regions with whole genome background. MSigDB pathway [96,97] was noted for genomic region enrichment analysis.

### GERP analysis

GERP (Genomic Evolutionary Rate Profiling) was performed to assess if highly TF bound cis-regulatory sites, categorized into promoter and enhancer-associated, correlates with increased evolutionary constraints. Highly constrained elements bed file containing high confidence regions (significant p-value) generated from per base GERP scores was retrieved from Sidow lab (http://mendel.stanford.edu/SidowLab/downloads/gerp/). Fraction of overlapping bases for each bins of “TF bound category” (low to high) with highly constrained elements was computed using bedtools-2.26.0 [94] and pandas-0.20.3, python2.7, further normalized by the fraction of “highly constrained elements” overlapping per 100 bp sized-region of TF bound categories. Additionally, Kolmogorov-Smirnov (KS) test was performed to evaluate statistically significant differences in distribution between the highly bound (20+ TF bound) and lowly bound regions (1-19 TF bound sites) for both promoter- and enhancer-associated sites.

### Co-binding analysis

Pairwise overlap of binding sites between each of the 208 TFs was performed with 50 bp up and downstream from the summit of peaks using python based pybedtools [93,94]. All other computations, and the pairwise peak overlap percentage for each TF to build the pairwise matrix, were performed using pandas-0.20.3, python2.7 [Python Software Foundation] to construct network plots, using R igraph, implementing Fruchterman Reingold algorithm. The interconnection between TF shared binding sites for 208 TFs was built with a minimum threshold of 75% or more overlap between any 2 factors. The sizes of vertices and nodes in the graph are representative of the number of connections each TF has with its connected partner, while edges represent the degree of overlap between TFs.

Co-binding was characterized by merging IDR-passing narrow peak files from 208 TFs with the “merge” function from the bedtools software package [98]. A minimum of 1 bp overlap was required and resultant peaks greater than 2 kb (~1%) were filtered from downstream analysis. Hierarchical clustering, using the Euclidean distance metric and Ward clustering method, of TFs based on degree of co-binding was performed in R with the “heatmap.2” function of the gplots package.

### LS-GKM SVM analysis

At peak level, LS-GKM support vector machines (SVMs) [99] were trained on a random sample of up to 5,000 narrow peaks (using all peaks for those with fewer) as a positive set against 10X random/null sequence with matched size, GC-content and repeat fraction as a negative set. At motif level, LS-GKM support vector machines (SVMs) [99] were trained on a sample of 5,000 random motif sites found by FIMO (MEME-suite), extending +/- 15 bp, for all DNA binding factors (n=171), as a positive set against the 10X random-null sequence with GC content and repeat fraction matched sequence as a negative set.

Null genomic sequences matched to observed binding events were obtained using the “nullseq_generate.py” function available with the LS-GKM package. The fold number of sequences (-x) was set to ten and the random seed (-r) was set to 1. SVMs were trained using the “gkmtrain” function with a kmer length (-l) of 11, kernel function (-t) of 4, regularization parameter (-c) of 1, number of informative columns (-k) of 7, and maximum number of mismatches (-d) of 3. Precision-recall area under the curves (PRAUC) were calculated by obtaining the 10-fold cross-validation results from “gkmtrain” (after setting the –x flag to 10), and inputting the results into the “pr.curve” function of the PRROC R package, resulting in mean PR-AUC of 0.66 at the peak level, and 0.74 at the motif level. Classifier values for all bound sequences were obtained using the “gkmpredict” function, and HOT sites (n=5,676) were scored with each DNA associated factor to assess their putative binding affinity at HOT regions, and percentile ranked to obtain top 5 percent and bottom 75 percent k-mer compared to enhancers with 2-10 associated TFs (n=5,676) and to random enhancers with any number of associated factors (0+) (n=5,676).

### Random Forest and PCA analysis

Principal Component Analysis (PCA) was performed on a DAP binding matrix composed of the presence or absence of motif in merged peaks as a binary matrix of loci, and implementing the python based ML library scikit-learn Sklearn (0.19.0) [100]. Plots for motif-based analyses were generated using the R package ggplot2 [101] and complex Heatmap [102]. Random Forest Classifier was trained on merged DAP binding matrices at both motif and peak level to predict cis-regulatory elements (promoter or enhancer, by IDEAS annotation) using the R package ranger [103], a faster implementation of random forest in R, and also tested using Sklearn 0.19.0. Median OOB (Out-of-bag) error estimate was computed for 100 instances of randomly sampled (n=1000) loci iterations, to compute the element classification and misclassification accuracy using confusion matrix.

### IP-mass spectrometry

Whole cell lysates of FLAG-tagged or unedited HepG2 cells (~20 million) were immunoprecipitated using a primary antibody raised against FLAG or the transcription factor, respectively. The IP fraction was loaded on a 12% TGX™ gel and separated with the Mini-PROTEAN^®^ Tetra Cell System (Bio-Rad). The whole lane was excised and sent to the University of Alabama at Birmingham Cancer Center Mass Spectrometry/Proteomics Shared Facility. The sample was analyzed on a LTQ XL Linear Ion Trap Mass Spectrometer by LC-ESI-MS/MS. Peptides were identified using SEQUEST tandem mass spectral analysis with probability based matching at p < 0.05. SEQUEST results were reported with ProteinProphet protXML Viewer (TPP v4.4 JETSTREAM) and filtered for a minimum probability of 0.9. For ENCODE Antibody Characterization Standards, all protein hits that met these criteria were reported, including common contaminants. Fold enrichment for each protein reported was determined using a custom script based on the FC-B score calculation [104]. Following ENCODE Antibody Characterization Guidelines, the transcription factor must be in the top 20 enriched proteins identified by IP-MS, and the top transcription factor overall for release. For GATAD2A co-associated TFs, the peptides with minimum 0.9 probability were present in less quantities than those of GATAD2A.

### Transcription factors footprints analysis

To identify TF footprints for comparison to ChIP-seq binding sites, we used PIQ (Protein Interaction Quantification) [86]. ENCODE HepG2 DNAse-seq raw FASTQs (paired-end 36 bp) of roughly equivalent size (Accession Numbers: ENCFF002EQ-G,H,I,J,M,N,O,P) were downloaded from the ENCODE portal and processed using ENCODE DNAse-seq standard pipeline (available at https://github.com/kundajelab/atac_dnase_pipelines) with flags: -species hg19 -nth 32 -memory 250G -dnase_seq -auto_detect_adapter -nreads 15000000 -ENCODE3. Processed BAM files were merged and used as input for PIQ TF footprinting using each TF’s top motif PWM. Next, identified TF footprints from every TF meeting a specified PIQ Purity (positive predictive value) were intersected with all identified ChIP-seq binding sites using BEDtools to correlate the number of unique TF footprints with the number of ChIP-seq factors identified at a given ChIP-seq binding site.

### SOM analysis

The self-organizing map was trained with the SOMatic package [67] using the previous chromatin analysis partitioning strategy [66] with modifications as described below We calculated the RPKM of each dataset’s first replicate over each of the 951,022 genomic segments to build a training matrix. We used each dataset’s second replicate to build a separate scoring matrix. The training matrix was used to train 5 trial self-organizing maps with a toroid topology with size 40 by 60 units using 10 million time steps (~10 epochs) and selected the best, based on fitting error using the scoring matrix, for further analysis, and segments were assigned to their closest units based on the scoring matrix.

To properly fit the data, SOM units with similar profiles across experiments were grouped into metaclusters using SOMatic. Briefly, metaclustering was performed using k-means clustering of the unit profiles to determine centroids for groups of units. Metaclusters were built around these centroids so that all of the units in a cluster remain connected. SOMatic’s metaclustering function attempts all metacluster numbers within a range given and scores them based on Akaike information criterion (AIC) [105]. The penalty term for this score is calculated using a parameter called the “dimensionality,” which is the number of independent dimensions in the data, which in this case are the individual cell subtypes. To estimate this number, we used a 60% cut on a hierarchical clustering done on the SOM unit vectors. For this work, the dimensionality was calculated to be 6. For metaclustering, all k between 50 and 250, with 64 trials, was tested and metacluster number 196 had the lowest AIC score and was chosen for further analysis.

To generate decision trees for these metaclusters, each of the segments in the training matrix was labeled with its final metacluster. For each metacluster, if the metacluster is of size n, n segments of other clusters were chosen randomly, and this set of positive and negative examples was split, using 80% of the examples for training and 20% for scoring. The training data was fed through an R script using the rpart and rattle packages to create, score, prune, and re-score a tree for each metacluster. This entire process was repeated for 100 trials with only the tree with the highest accuracy drawn.

## Acknowledgements

Research reported in this publication was supported by the National Human Genome Research Institute of the National Institutes of Health under Award Number U54HG006998 to R.M.M. and E.M.M. The content is solely the responsibility of the authors and does not necessarily represent the official views of the National Institutes of Health. This work was also supported by funds from The HudsonAlpha Institute for Biotechnology. We thank Rosy Nguyen, Dianna Moore, and Megan McEown for their outstanding technical efforts in this study. We thank Brian S. Roberts and Gregory M. Cooper for helpful comments, HudsonAlpha’s Genomic Services Laboratory led by Dr. Shawn Levy for the high-throughput sequencing of much of the data used in this paper, and members of the ENCODE Consortium for public deposition of data generated by other Consortium groups.

## References

1 Vaquerizas JM, Kummerfeld SK, Teichmann SA, Luscombe NM. 2009. A census of human transcription factors: function, expression and evolution. Nat Rev Genet 10: 252–63.

2 Lambert SA, Jolma A, Campitelli LF, Das PK, et al. 2018. The Human Transcription Factors. Cell 172: 650–65.

3 Wingender E, Schoeps T, Donitz J. 2013. TFClass: an expandable hierarchical classification of human transcription factors. Nucleic Acids Res 41: D165–70.

4 Weirauch MT, Yang A, Albu M, Cote AG, et al. 2014. Determination and inference of eukaryotic transcription factor sequence specificity. Cell 158: 1431–43.

5 Cowper-Sallari R, Zhang X, Wright JB, Bailey SD, et al. 2012. Breast cancer risk-associated SNPs modulate the affinity of chromatin for FOXA1 and alter gene expression. Nat Genet 44: 1191–8.

6 Dror I, Golan T, Levy C, Rohs R, et al. 2015. A widespread role of the motif environment in transcription factor binding across diverse protein families. Genome Res 25: 1268–80.

7 Vernimmen D, Bickmore WA. 2015. The Hierarchy of Transcriptional Activation: From Enhancer to Promoter. Trends Genet 31: 696–708.

8 Yosef N, Shalek AK, Gaublomme JT, Jin H, et al. 2013. Dynamic regulatory network controlling TH17 cell differentiation. Nature 496: 461–8.

9 Dasen JS, Tice BC, Brenner-Morton S, Jessell TM. 2005. A Hox regulatory network establishes motor neuron pool identity and target-muscle connectivity. Cell 123: 477–91.

10 Busskamp V, Lewis NE, Guye P, Ng AH, et al. 2014. Rapid neurogenesis through transcriptional activation in human stem cells. Mol Syst Biol 10: 760.

11 Chen X, Xu H, Yuan P, Fang F, et al. 2008. Integration of external signaling pathways with the core transcriptional network in embryonic stem cells. Cell 133: 1106–17.

12 Black JB, Adler AF, Wang HG, D’Ippolito AM, et al. 2016. Targeted Epigenetic Remodeling of Endogenous Loci by CRISPR/Cas9-Based Transcriptional Activators Directly Converts Fibroblasts to Neuronal Cells. Cell Stem Cell 19: 406–14.

13 Visel A, Blow MJ, Li Z, Zhang T, et al. 2009. ChIP-seq accurately predicts tissue-specific activity of enhancers. Nature 457: 854–8.

14 Iwafuchi-Doi M, Zaret KS. 2014. Pioneer transcription factors in cell reprogramming. Genes Dev 28: 2679–92.

15 Johnson DS, Mortazavi A, Myers RM, Wold B. 2007. Genome-wide mapping of in vivo protein-DNA interactions. Science 316: 1497–502.

16 Mikkelsen TS, Ku M, Jaffe DB, Issac B, et al. 2007. Genome-wide maps of chromatin state in pluripotent and lineage-committed cells. Nature 448: 553–60.

17 Robertson G, Hirst M, Bainbridge M, Bilenky M, et al. 2007. Genome-wide profiles of STAT1 DNA association using chromatin immunoprecipitation and massively parallel sequencing. Nat Methods 4: 651–7.

18 Lambert SA, Albu M, Hughes TR, Najafabadi HS. 2016. Motif comparison based on similarity of binding affinity profiles. Bioinformatics 32: 3504–6.

19 Najafabadi HS, Albu M, Hughes TR. 2015. Identification of C2H2-ZF binding preferences from ChIP-seq data using RCADE. Bioinformatics 31: 2879–81.

20 Bailey TL, Boden M, Buske FA, Frith M, et al. 2009. MEME SUITE: tools for motif discovery and searching. Nucleic Acids Res 37: W202–8.

21 Bailey TL, Machanick P. 2012. Inferring direct DNA binding from ChIP-seq. Nucleic Acids Res 40: e128.

22 Landolin JM, Johnson DS, Trinklein ND, Aldred SF, et al. 2010. Sequence features that drive human promoter function and tissue specificity. Genome Res 20: 890–8.

23 Whitfield TW, Wang J, Collins PJ, Partridge EC, et al. 2012. Functional analysis of transcription factor binding sites in human promoters. Genome Biol 13: R50.

24 Hallikas O, Palin K, Sinjushina N, Rautiainen R, et al. 2006. Genome-wide prediction of mammalian enhancers based on analysis of transcription-factor binding affinity. Cell 124: 47–59.

25 Levo M, Zalckvar E, Sharon E, Dantas Machado AC, et al. 2015. Unraveling determinants of transcription factor binding outside the core binding site. Genome Res 25: 1018–29.

26 Garton M, Najafabadi HS, Schmitges FW, Radovani E, et al. 2015. A structural approach reveals how neighbouring C2H2 zinc fingers influence DNA binding specificity. Nucleic Acids Res 43: 9147–57.

27 Hauser K, Essuman B, He Y, Coutsias E, et al. 2016. A human transcription factor in search mode. Nucleic Acids Res 44: 63–74.

28 Slattery M, Riley T, Liu P, Abe N, et al. 2011. Cofactor binding evokes latent differences in DNA binding specificity between Hox proteins. Cell 147: 1270–82.

29 Siggers T, Reddy J, Barron B, Bulyk ML. 2014. Diversification of transcription factor paralogs via noncanonical modularity in C2H2 zinc finger DNA binding. Mol Cell 55: 640–8.

30 Siggers T, Gordan R. 2014. Protein-DNA binding: complexities and multi-protein codes. Nucleic Acids Res 42: 2099–111.

31 Gertz J, Savic D, Varley KE, Partridge EC, et al. 2013. Distinct properties of cell-type-specific and shared transcription factor binding sites. Mol Cell 52: 25–36.

32 Reddy TE, Pauli F, Sprouse RO, Neff NF, et al. 2009. Genomic determination of the glucocorticoid response reveals unexpected mechanisms of gene regulation. Genome Res 19: 2163–71.

33 Chen X, Yu B, Carriero N, Silva C, et al. 2017. Mocap: large-scale inference of transcription factor binding sites from chromatin accessibility. Nucleic Acids Res 45: 4315–29.

34 Garber M, Yosef N, Goren A, Raychowdhury R, et al. 2012. A high-throughput chromatin immunoprecipitation approach reveals principles of dynamic gene regulation in mammals. Mol Cell 47: 810–22.

35 Wang J, Zhuang J, Iyer S, Lin X, et al. 2012. Sequence features and chromatin structure around the genomic regions bound by 119 human transcription factors. Genome Res 22: 1798–812.

36 Schmitges FW, Radovani E, Najafabadi HS, Barazandeh M, et al. 2016. Multiparameter functional diversity of human C2H2 zinc finger proteins. Genome Res 26: 1742–52.

37 Imbeault M, Helleboid PY, Trono D. 2017. KRAB zinc-finger proteins contribute to the evolution of gene regulatory networks. Nature 543: 550–4.

38 Yan J, Enge M, Whitington T, Dave K, et al. 2013. Transcription factor binding in human cells occurs in dense clusters formed around cohesin anchor sites. Cell 154: 801–13.

39 Savic D, Partridge EC, Newberry KM, Smith SB, et al. 2015. CETCh-seq: CRISPR epitope tagging ChIP-seq of DNA-binding proteins. Genome Res 25: 1581–9.

40 Partridge EC, Watkins TA, Mendenhall EM. 2016. Every transcription factor deserves its map: Scaling up epitope tagging of proteins to bypass antibody problems. Bioessays 38: 801–11.

41 Baresic M, Salatino S, Kupr B, van Nimwegen E, et al. 2014. Transcriptional network analysis in muscle reveals AP-1 as a partner of PGC-1alpha in the regulation of the hypoxic gene program. Mol Cell Biol 34: 2996–3012.

42 Fernandez PC, Frank SR, Wang L, Schroeder M, et al. 2003. Genomic targets of the human c-Myc protein. Genes Dev 17: 1115–29.

43 Landt SG, Marinov GK, Kundaje A, Kheradpour P, et al. 2012. ChIP-seq guidelines and practices of the ENCODE and modENCODE consortia. Genome Res 22: 1813–31.

44 Baranello L, Kouzine F, Sanford S, Levens D. 2016. ChIP bias as a function of cross-linking time. Chromosome Res 24: 175–81.

45 Teytelman L, Ozaydin B, Zill O, Lefrancois P, et al. 2009. Impact of chromatin structures on DNA processing for genomic analyses. PLoS One 4: e6700.

46 Kharchenko PV, Tolstorukov MY, Park PJ. 2008. Design and analysis of ChIP-seq experiments for DNA-binding proteins. Nat Biotechnol 26: 1351–9.

47 Li Q, Brown JB, Huang H, Bickel PJ. 2011. Measuring reproducibility of high-throughput experiments. The Annals of Applied Statistics 5: 1752–79.

48 Zhang Y, An L, Yue F, Hardison RC. 2016. Jointly characterizing epigenetic dynamics across multiple human cell types. Nucleic Acids Res 44: 6721–31.

49 Kowalczyk MS, Hughes JR, Garrick D, Lynch MD, et al. 2012. Intragenic enhancers act as alternative promoters. Mol Cell 45: 447–58.

50 Dao LTM, Galindo-Albarran AO, Castro-Mondragon JA, Andrieu-Soler C, et al. 2017. Genome-wide characterization of mammalian promoters with distal enhancer functions. Nat Genet 49: 1073–81.

51 Andersson R, Sandelin A, Danko CG. 2015. A unified architecture of transcriptional regulatory elements. Trends Genet 31: 426–33.

52 Cirillo LA, Lin FR, Cuesta I, Friedman D, et al. 2002. Opening of compacted chromatin by early developmental transcription factors HNF3 (FoxA) and GATA-4. Mol Cell 9: 279–89.

53 Magnani L, Eeckhoute J, Lupien M. 2011. Pioneer factors: directing transcriptional regulators within the chromatin environment. Trends Genet 27: 465–74.

54 Teytelman L, Thurtle DM, Rine J, van Oudenaarden A. 2013. Highly expressed loci are vulnerable to misleading ChIP localization of multiple unrelated proteins. Proc Natl Acad Sci U S A 110: 18602–7.

55 Bailey TL, Johnson J, Grant CE, Noble WS. 2015. The MEME Suite. Nucleic Acids Res 43: W39–49.

56 Machanick P, Bailey TL. 2011. MEME-ChIP: motif analysis of large DNA datasets. Bioinformatics 27: 1696–7.

57 Ma W, Noble WS, Bailey TL. 2014. Motif-based analysis of large nucleotide data sets using MEME-ChIP. Nat Protoc 9: 1428–50.

58 Gupta S, Stamatoyannopoulos JA, Bailey TL, Noble WS. 2007. Quantifying similarity between motifs. Genome Biol 8: R24.

59 Sandelin A, Alkema W, Engstrom P, Wasserman WW, et al. 2004. JASPAR: an open-access database for eukaryotic transcription factor binding profiles. Nucleic Acids Res 32: D91–4.

60 Mathelier A, Fornes O, Arenillas DJ, Chen CY, et al. 2016. JASPAR 2016: a major expansion and update of the open-access database of transcription factor binding profiles. Nucleic Acids Res 44: D110–5.

61 Oliphant AR, Brandl CJ, Struhl K. 1989. Defining the sequence specificity of DNA-binding proteins by selecting binding sites from random-sequence oligonucleotides: analysis of yeast GCN4 protein. Mol Cell Biol 9: 2944–9.

62 Worsley Hunt R, Wasserman WW. 2014. Non-targeted transcription factors motifs are a systemic component of ChIP-seq datasets. Genome Biol 15: 412.

63 Jolma A, Yan J, Whitington T, Toivonen J, et al. 2013. DNA-binding specificities of human transcription factors. Cell 152: 327–39.

64 Morgunova E, Taipale J. 2017. Structural perspective of cooperative transcription factor binding. Curr Opin Struct Biol 47: 1–8.

65 Wei B, Jolma A, Sahu B, Orre LM, et al. 2018. A protein activity assay to measure global transcription factor activity reveals determinants of chromatin accessibility. Nat Biotechnol 36: 521–9.

66 Mortazavi A, Pepke S, Jansen C, Marinov GK, et al. 2013. Integrating and mining the chromatin landscape of cell-type specificity using self-organizing maps. Genome Res 23: 2136–48.

67 Longabaugh WJR, Zeng W, Zhang JA, Hosokawa H, et al. 2017. Bcl11b and combinatorial resolution of cell fate in the T-cell gene regulatory network. Proc Natl Acad Sci U S A 114: 5800–7.

68 McLean CY, Bristor D, Hiller M, Clarke SL, et al. 2010. GREAT improves functional interpretation of cis-regulatory regions. Nat Biotechnol 28: 495–501.

69 Whyte WA, Bilodeau S, Orlando DA, Hoke HA, et al. 2012. Enhancer decommissioning by LSD1 during embryonic stem cell differentiation. Nature 482: 221–5.

70 Liang Z, Brown KE, Carroll T, Taylor B, et al. 2017. A high-resolution map of transcriptional repression. Elife 6.

71 Zhang Y, Ng HH, Erdjument-Bromage H, Tempst P, et al. 1999. Analysis of the NuRD subunits reveals a histone deacetylase core complex and a connection with DNA methylation. Genes Dev 13: 1924–35.

72 Huttlin EL, Ting L, Bruckner RJ, Gebreab F, et al. 2015. The BioPlex Network: A Systematic Exploration of the Human Interactome. Cell 162: 425–40.

73 Faherty N, Benson M, Sharma E, Lee A, et al. 2016. Negative autoregulation of BMP dependent transcription by SIN3B splicing reveals a role for RBM39. Sci Rep 6: 28210.

74 Choi WI, Yoon JH, Kim MY, Koh DI, et al. 2014. Promyelocytic leukemia zinc finger-retinoic acid receptor alpha (PLZF-RARalpha), an oncogenic transcriptional repressor of cyclin-dependent kinase inhibitor 1A (p21WAF/CDKN1A) and tumor protein p53 (TP53) genes. J Biol Chem 289: 18641–56.

75 Hnisz D, Abraham BJ, Lee TI, Lau A, et al. 2013. Super-enhancers in the control of cell identity and disease. Cell 155: 934–47.

76 Gunther K, Rust M, Leers J, Boettger T, et al. 2013. Differential roles for MBD2 and MBD3 at methylated CpG islands, active promoters and binding to exon sequences. Nucleic Acids Res 41: 3010–21.

77 Zaret KS, Carroll JS. 2011. Pioneer transcription factors: establishing competence for gene expression. Genes Dev 25: 2227–41.

78 Lian Z, Karpikov A, Lian J, Mahajan MC, et al. 2008. A genomic analysis of RNA polymerase II modification and chromatin architecture related to 3’ end RNA polyadenylation. Genome Res 18: 1224–37.

79 Conacci-Sorrell M, McFerrin L, Eisenman RN. 2014. An overview of MYC and its interactome. Cold Spring Harb Perspect Med 4: a014357.

80 Hervouet E, Vallette FM, Cartron PF. 2009. Dnmt3/transcription factor interactions as crucial players in targeted DNA methylation. Epigenetics 4: 487–99.

81 Boyle AP, Araya CL, Brdlik C, Cayting P, et al. 2014. Comparative analysis of regulatory information and circuits across distant species. Nature 512: 453–6.

82 Gerstein MB, Lu ZJ, Van Nostrand EL, Cheng C, et al. 2010. Integrative analysis of the Caenorhabditis elegans genome by the modENCODE project. Science 330: 1775–87.

83 Moorman C, Sun LV, Wang J, de Wit E, et al. 2006. Hotspots of transcription factor colocalization in the genome of Drosophila melanogaster. Proc Natl Acad Sci U S A 103: 12027–32.

84 Wreczycka K, Franke V, Uyar B, Wurmus R, et al. 2017. HOT or not: Examining the basis of high-occupancy target regions. bioRxiv doi: 101101/107680.

85 Shin H, Liu T, Duan X, Zhang Y, et al. 2013. Computational methodology for ChIP-seq analysis. Quant Biol 1: 54–70.

86 Sherwood RI, Hashimoto T, O’Donnell CW, Lewis S, et al. 2014. Discovery of directional and nondirectional pioneer transcription factors by modeling DNase profile magnitude and shape. Nat Biotechnol 32: 171–8.

87 Panne D, Maniatis T, Harrison SC. 2007. An atomic model of the interferon-beta enhanceosome. Cell 129: 1111–23.

88 Li H, Durbin R. 2009. Fast and accurate short read alignment with Burrows-Wheeler transform. Bioinformatics 25: 1754–60.

89 Li H, Handsaker B, Wysoker A, Fennell T, et al. 2009. The Sequence Alignment/Map format and SAMtools. Bioinformatics 25: 2078–9.

90 Worsley Hunt R, Mathelier A, Del Peso L, Wasserman WW. 2014. Improving analysis of transcription factor binding sites within ChIP-Seq data based on topological motif enrichment. BMC Genomics 15: 472.

91 Teng M, Irizarry RA. 2017. Accounting for GC-content bias reduces systematic errors and batch effects in ChIP-seq data. Genome Res 27: 1930–8.

92 Uhlen M, Fagerberg L, Hallstrom BM, Lindskog C, et al. 2015. Proteomics. Tissue-based map of the human proteome. Science 347: 1260419.

93 Dale RK, Pedersen BS, Quinlan AR. 2011. Pybedtools: a flexible Python library for manipulating genomic datasets and annotations. Bioinformatics 27: 3423–4.

94 Quinlan AR, Hall IM. 2010. BEDTools: a flexible suite of utilities for comparing genomic features. Bioinformatics 26: 841–2.

95 Fletez-Brant C, Lee D, McCallion AS, Beer MA. 2013. kmer-SVM: a web server for identifying predictive regulatory sequence features in genomic data sets. Nucleic Acids Res 41: W544–56. 96

96 Liberzon A, Birger C, Thorvaldsdottir H, Ghandi M, et al. 2015. The Molecular Signatures Database (MSigDB) hallmark gene set collection. Cell Syst 1: 417–25.

97 Subramanian A, Tamayo P, Mootha VK, Mukherjee S, et al. 2005. Gene set enrichment analysis: a knowledge-based approach for interpreting genome-wide expression profiles. Proc Natl Acad Sci U S A 102: 15545–50.

98 Quinlan AR. 2014. BEDTools: The Swiss-Army Tool for Genome Feature Analysis. Curr Protoc Bioinformatics 47: 112 1–34.

99 Ghandi M, Mohammad-Noori M, Ghareghani N, Lee D, et al. 2016. gkmSVM: an R package for gapped-kmer SVM. Bioinformatics 32: 2205–7.

100 Pedregosa ea. 2011. Scikit-learn: Machine Learning in Python. JMLR 12: 2825–30.

101 Wickham H. 2016. ggplot2: Elegant Graphics for Data Analysis. Springer-Verlag, New York.

102 Gu Z, Eils R, Schlesner M. 2016. Complex heatmaps reveal patterns and correlations in multidimensional genomic data. Bioinformatics 32: 2847–9.

103 Wright MN, Ziegler, A. 2017. ranger: A fast implementation of random forests for high dimensional data in C++ and R. Journal of Statistical Software 77: 1–17.

104 Mellacheruvu D, Wright Z, Couzens AL, Lambert JP, et al. 2013. The CRAPome: a contaminant repository for affinity purification-mass spectrometry data. Nat Methods 10: 730–6.

105 Akaike H. 1973. Information theory and an extension of the maximum likelihood principle. International Symposium on Information Theory: 267–81.

